# Magnetic resonance spectroscopy analysis of intramyocellular lipid composition in lipodystrophic patients and athletes

**DOI:** 10.1101/382283

**Authors:** David B Savage, Laura Watson, Katie Carr, Claire Adams, Soren Brage, Krishna K Chatterjee, Leanne Hodson, Chris Boesch, Graham J Kemp, Alison Sleigh

**Author notes:** Correspondence and Reprint Requests: A. Sleigh, Box 65 Addenbrooke’s Hospital, Hills Road, Cambridge, CB2 0QQ, United Kingdom. Disclosure Summary: The authors have nothing to disclose.

## Abstract

**Context:** Paradoxically, intramyocellular lipid (IMCL) accumulation has been linked to both insulin-resistant and to insulin-sensitive (athletes) states. The composition of this lipid store is unknown in these states.

**Design and Methods:** We used a recently validated and potentially widely applicable ^1^H magnetic resonance spectroscopy method to compare the compositional saturation index (CH_2_:CH_3_ ratio) and concentration independent of composition (CH_3_) of intramyocellular lipid in the soleus and tibialis anterior muscles of 16 female insulin-resistant lipodystrophic patients with that of age- and gender-matched athletes (n=14) and healthy controls (n = 41).

**Main Outcome:** IMCL compositional saturation index (CH_2_:CH_3_ ratio).

**Results:** The IMCL CH_2_:CH_3_ ratio was significantly higher in both muscles of the lipodystrophic patients compared with age- and gender-matched controls but not compared to athletes. IMCL CH_2_:CH_3_ was dependent on IMCL concentration in the controls and after adjusting the composition index for quantity (CH_2_:CH_3adj_) was able to distinguish patients from athletes. With groups pooled, this CH_2_:CH_3adj_ marker had the strongest relation to insulin resistance (HOMA-IR) compared to other measures of lipid concentration and composition, especially in the soleus muscle. Contrary to the ‘athlete’s paradox’, IMCL in athletes was similar in tibialis anterior (p>0.05) and significantly lower in the soleus (p < 0.004) compared to both controls and patients.

**Conclusions:** The IMCL saturation index adjusted for quantity, which likely reflects accumulation of saturated IMCL, is more closely associated with insulin resistance than concentration alone.

## Introduction

Following the demonstration that ^1^H magnetic resonance spectroscopy (^1^H MRS) can noninvasively distinguish intramyocellular (IMCL) from extramyocellular (EMCL) lipids (1,2), associations were reported between soleus IMCL accumulation and insulin resistance (IR) independent of fat mass (3–5). Given that skeletal muscle accounts for a significant amount of insulin-stimulated glucose disposal and thus represents the primary site for postprandial glucose disposal (6), these findings were of considerable physiological interest. Furthermore these data strongly supported the link between ectopic fat accumulation and insulin resistance (7,8). Although it soon became clear that triglycerides themselves were unlikely to be involved in causing insulin resistance, intramuscular triglyceride content does appear to correlate with insulin resistance in some states (5,9–12). One particularly striking and surprising exception to this was reported in athletes, where histological studies suggested that neutral lipid accumulation was a feature of skeletal muscle in insulin-sensitive trained athletes (13,14), and this has led to the now widely cited notion of an ‘athlete’s paradox’ (13). This concept is consistent with the idea that triglyceride itself is not involved in causing insulin resistance and has prompted several efforts to identify the lipid intermediates responsible for causing insulin resistance or preserving the insulin sensitivity of athletes.

Saturated fat has been implicated in the pathogenesis of metabolic disease (15,16) and we have recently described and validated (using IMCL/EMCL simulated phantoms of known composition) a ^1^H MRS method that provides an *in vivo* compositional marker of IMCL that primarily reflects the degree of saturation of the fatty acid chains within triglyceride (17). This marker, which we call the ‘IMCL saturation index’ (CH_2_:CH_3_), utilizes good quality spectra acquired at 3T with short echo time and compares the CH_2_ resonance located at 1.3 ppm (which is influenced by both concentration and composition), with that of the CH_3_ resonance at 0.9 ppm which is independent of triglyceride composition: this is illustrated in Figure 1. From this figure it can also be seen that using concentration of ‘H’ that resonate at 1.3 ppm (CH_2_) to represent the concentration of lipid without knowledge of the underlying composition, as has been the practice in virtually all published ^1^H MRS studies of IMCL so far, confounds the contributions of both the concentration of lipid and its composition. This has potential for significant error in the estimation of concentration: the composition could contribute as much as 43% to the observed signal, which is equivalent to a 77 *%* theoretical increase in signal if the pool was stearic acid instead of sunflower oil. We therefore use the IMCL CH_3_ peak at 0.9 ppm to estimate the total concentration of IMCL, as this is independent of the degree of saturation of the fatty acid chains within triglyceride (i.e. composition). We call this the *composition-independent* IMCL concentration estimate, to distinguish it from the *conventional* estimate using CH_2_.

**Figure 1.**
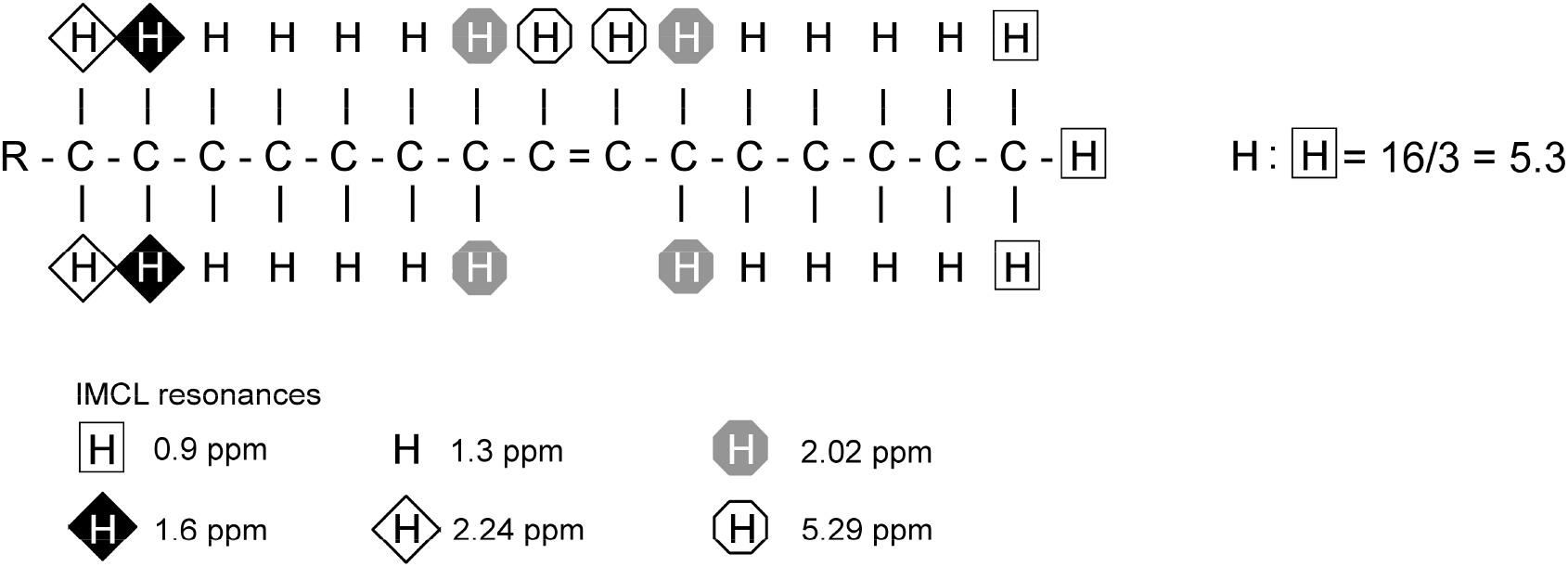
The ratio of CH_2_:CH_3_ is influenced primarily by the degree of saturation of the fatty acids within triglyceride. This figure shows the principle of the CH_2_:CH_3_ ratio as a compositional saturation index of IMCL. In this example palmitoleic acid component of TG has a theoretical ratio of CH_2_ (at 1.3 ppm) to CH_3_ (at 0.9 ppm) of 16/3 = 5.3 which is lower than the equivalent ratio of palmitic acid = 8.0. This is due not only to the desaturation of two CH_2_ but also to the alteration in the chemical environment of the neighbouring CH_2_. For FA chains of equal number of double bonds, the CH_2_:CH_3_ ratio will also be scaled by chain length, although this will have a proportionally smaller effect (17).

Lipodystrophy is a rare cause of severe IR and is typically characterized by prominent ectopic fat accumulation due both to the reduction in adipocyte lipid storage capacity and the associated hyperphagia induced by leptin deficiency. In order to ascertain if intramyocellular lipid composition is altered in lipodystrophic (LD) patients, and if such changes in lipid composition might help to elucidate the ‘athlete’s paradox’, we determined the compositional saturation index (CH_2_:CH_3_ ratio) and composition-independent concentration (from CH_3_) of IMCL in the soleus and tibialis anterior muscles of female insulin-resistant lipodystrophic patients, as well as age- and gender-matched athletes and non-athlete controls.

## Material and Methods

### Participants

Sixteen female patients with lipodystrophy were identified as part of a long-standing study of human IR syndromes, while age- and gender-matched controls (n=41) and athletes (n=14) were recruited by advertisement. Soleus IMCL data from five of the patients were included in a previously published study (18). Control and athlete exclusion criteria included smoking, drug or alcohol addiction, any current or past medical disorder or medications that could affect measurements including supplements such as creatine, and standard MR contraindications. Controls were recruited who exercised less than 3 times per week for 1 hour each time, whilst the athletes were part of a running club, some at international level, and all ran distances between 10 and 40 km regularly. The patient studies were approved by the National Health Service Research Ethics Committee, the healthy volunteer studies by East of England Cambridge Central Ethics Committee. Studies were conducted in accordance with the Declaration of Helsinki and all participants provided written informed consent.

### Protocol

Volunteers were instructed to follow normal dietary habits for 3 days prior to arrival at the Cambridge NIHR/Wellcome Trust Clinical Research Facility. Controls formed part of a larger study and arrived at 14:00 on Day 1 where a subset (24/41) performed a sub-maximal exercise test at 15:30, all participants were given an energy balanced standardised evening meal (30-35 % fat, 50-55 % carbohydrate, 12-15 % protein) at 18:30. On Day 2 fasting blood samples were taken and a light breakfast provided before ^1^H MRS at 09:30. Athletes were instructed to refrain from vigorous exercise for at least 24 hours prior to arrival at 08:00 when fasting blood samples were taken before participants consumed a light breakfast, before ^1^H MRS at 09:30 followed by a maximal VO_2max_ exercise test. Lipodystrophic patients also had ^1^H MRS in the fed state but did not perform an exercise test.

HOMA-IR was calculated as fasting insulin (pmol/l)*fasting glucose (μU/ml) / 22.5. Body composition was assessed by Dual-energy X-ray Absorptiometry (GE Lunar Prodigy encore v12.5 or GE Lunar iDXA encore v16 (athletes).

### ^1^H Magnetic Resonance Spectroscopy (^1^H MRS)

^1^H MRS studies were performed on a Siemens 3T scanner (Erlangen, Germany) using the Point-Resolved Spectroscopy (PRESS) sequence with short echo time of 35 ms. A water-suppressed ^1^H spectrum was acquired from a voxel of cube length 1.3 cm positioned to avoid visible fat on T_1_-weighted images within TA and SOL, using a 5 s repetition time and 64 averages (non-water-suppressed spectrum 4 averages). Data were analysed in jMRUI (19,20) and fitted with the AMARES (21) algorithm using identical prior knowledge parameters: Gaussian lineshapes (except water: Lorentzian), soft constraints on EMCL/IMCL CH_2_ frequencies and linewidths, CH_3_ resonant frequencies and linewidths determined from known and inferred prior knowledge relative to the CH_2_ resonance (22), and with all amplitudes estimated. IMCL CH_2_ and CH_3_ are quantified relative to the methyl group of creatine plus phosphocreatine at 3.0 ppm (Cr). As this resonance exhibits different lineshape characteristics in the tibialis anterior and soleus muscles (23), comparable quantification between muscles using a nominal concentration of muscle creatine is not valid; instead a scaling factor of Cr to water signal for each muscle was established from a subset of participants who had nonwater-suppressed datasets, yielding a calculated water signal (water-calc). Absolute composition-independent IMCL concentrations in mmol/kg muscle wet weight (ww) were calculated from the compositionally invariant CH_3_ IMCL resonance, with standard assumptions regarding muscle water content, and correction for T2 relaxation effects, J coupling and proton density as outlined in the supplementary information (24). The IMCL saturation index (CH_2_:CH_3_) was calculated as IMCL CH_2_/CH_3_, and the IMCL saturation index adjusted for quantity (CH_2_:CH_3adj_) = CH_2_ – (mCH_3_ + c), where m and c are the gradient and intercept of the regression line through the control data points of CH_2_ vs CH_3_. Investigators were blind to the insulin resistance status of the participants during ^1^H MRS analysis.

### Assessment of VO_2max_

Participants underwent continuous incremental exercise testing to 85% age-predicted maximum heart rate (controls) or volitional exhaustion (athletes) on a treadmill (Trackmaster TMX425, Med-electronics, MD). Oxygen consumption was measured using a spiro-ergometer (Medical Graphics UK Ltd and Breezesuit^®^ Gas exchange software). For the control participants a standard incremental protocol was performed (25), while the athletes undertook a protocol that began with a 10 min warm-up period at each participant’s preferred warm-up running speed, following which the test was initiated at 9 km/hr and increased steadily (0.74 km.hr^-1^.min^-1^), with a ramp at 5 minutes (increasing 0.5 % every 15s) until exhaustion or a plateaux in VO_2_ was apparent. In controls VO_2max_ was calculated by extrapolating the submaximal heart rate – VO_2_ relationship to age-predicted maximum heart rate (26).

### Statistics

All statistics were performed in IBM SPSS Statistics 24 (IBM, Armonk, NY: IBM Corp.) with significance set at P < 0.05. Normality was assessed by the Shapiro-Wilk test and non-normally distributed data were log-transformed prior to statistical testing. ANOVA with Games-Howell post hoc analysis was used to compare means between groups, and Pearson’s correlation coefficient for analyzing associations. Due to the non-normality of LN(HOMA-IR), IMCL associations with HOMA-IR were assessed by Spearman’s rank correlation coefficient. Data are mean ± SEM.

## Results

### Participants

Of the insulin-resistant patients with lipodystrophy 13 had partial forms (8 patients with FPLD2 due to LMNA mutations, 5 patients with FPLD3 due to PPARG mutations), and 3 generalized lipodystrophy (GLD, 2 patients with an acquired form – AGLD, and 1 due to mutations in the PCYT1A gene (27)). EMCL was absent in the 2 patients with AGLD (Figure 2), but those with partial forms had EMCL such that overall patients’ EMCL was similar to both controls and athletes (Table 1). The age- and gender-matched controls had a wide range of BMI (19.6 – 35.6 kg.m^-2^) and HOMA-IR (0.3 – 4.9). As a group, insulin and HOMA-IR were significantly higher in the LD patients and lower in the athletes (Table 1, Figure 3A) compared with controls, as expected. Fat mass and percentage body fat were similar between LD patients and athletes, which were both lower compared to controls (Table 1, Figure 3B). Serum triglycerides were higher whilst high-density lipoprotein (HDL)-cholesterol concentrations were lower in the LD patients (Table 1) compared to either controls or athletes.

**Figure 2.**
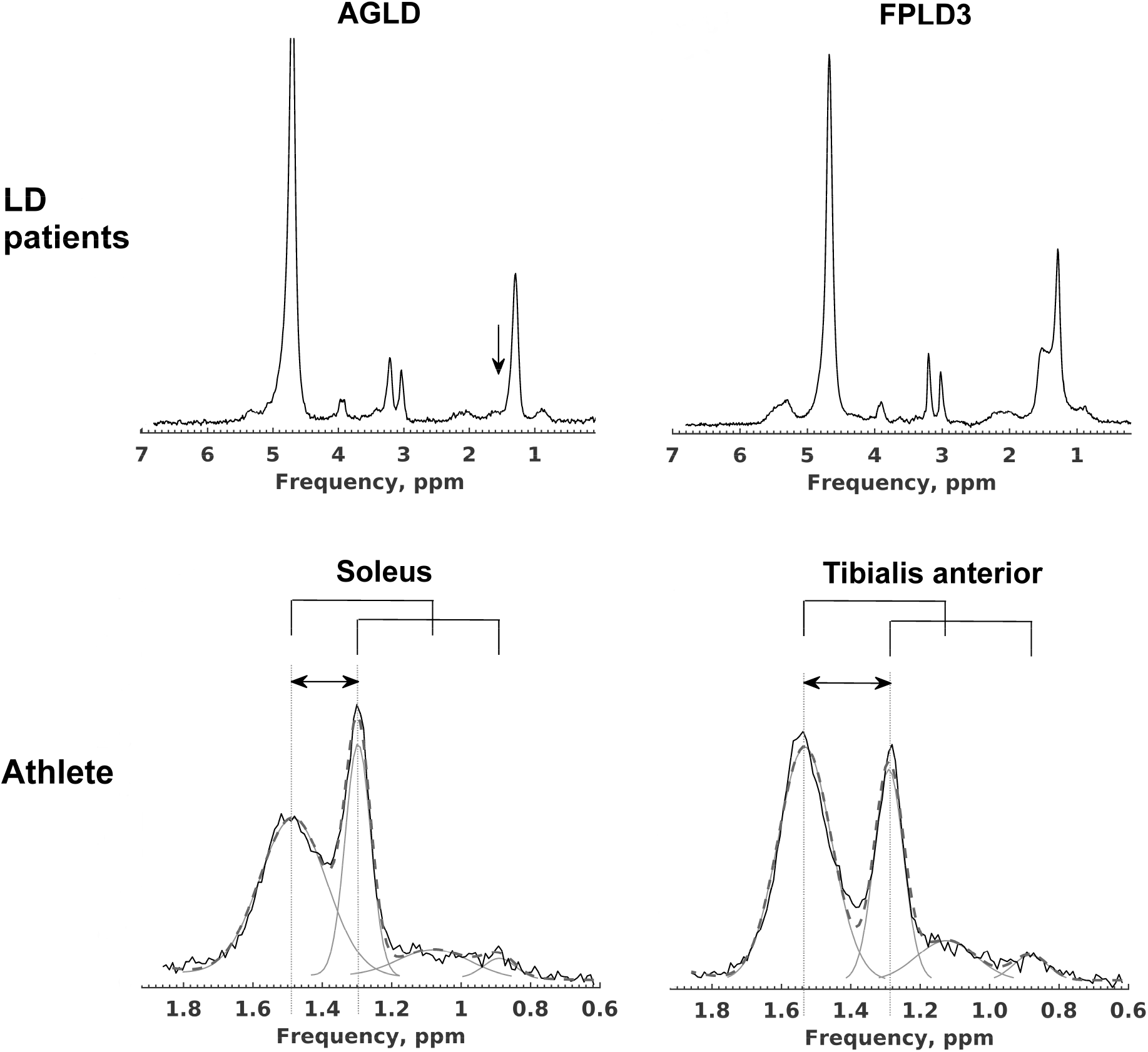
Representative ^1^H MRS spectra from lipodystrophic (LD) patients and an athlete. Water-suppressed spectra from the soleus muscle of a patient with acquired generalised lipodystrophy (AGLD, *upper left*) and familial partial lipodystrophy (FPLD3, *upper right*); the absence of extramyocellular lipid (EMCL) in AGLD is highlighted by the vertical arrow. Spectra from an athlete’s soleus (*lower left*) and tibialis anterior (*lower right*) muscles illustrating the raw data (solid black) and overall fit (grey dashed) and individual fit components (solid grey) in the frequency range that contains the EMCL CH_2_ (~ 1.5 ppm), CH_3_ (~1.1 ppm) and IMCL CH_2_ (1.3 ppm) and CH_3_ (0.9 ppm) resonances. This volunteer had the lowest soleus IMCL concentration of all participants (2.0 mmol/kg ww). The horizontal arrows highlights that the EMCL resonances are systematically shifted very slightly upfield in the soleus muscle compared with the tibialis anterior due to fibre orientation effects. The CH_3_ resonant frequencies are linked to the CH_2_ frequencies (solid bridge lines above spectra) and are also shifted. The fitting procedure fixes the relative CH_2_ to CH_3_ frequency shift for both EMCL and IMCL, and also fixes the CH_3_ linewidth relative to the corresponding CH_2_, but permits soft constraints on the EMCL frequencies.

**Figure 3.**
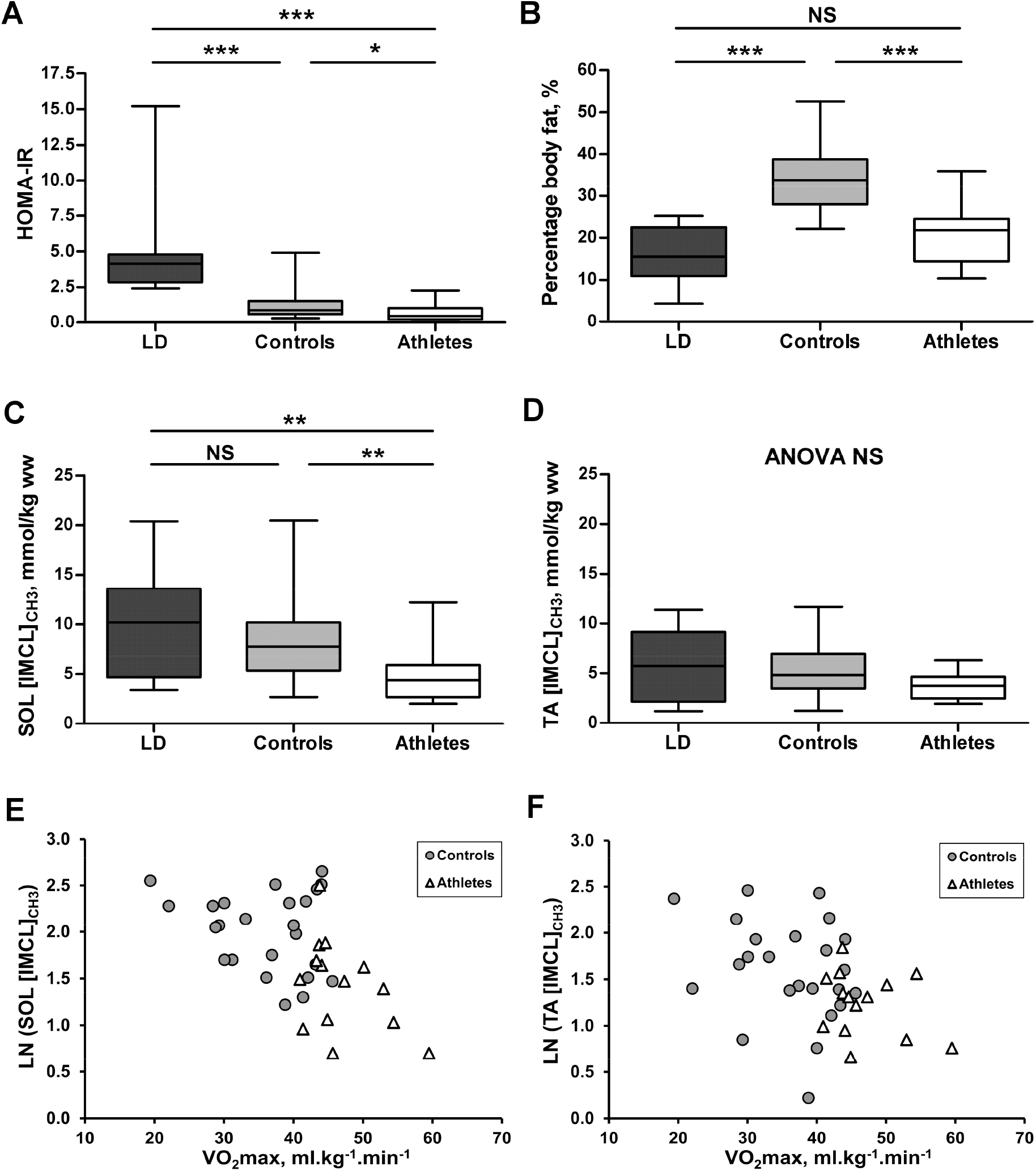
Participant characteristics and composition-independent concentrations of intramyocellular lipid in lipodystrophic patients (black), controls (grey), and athletes (white). (A-D) Box and whisker plots showing (A) Whole-body insulin sensitivity assessed by HOMA-IR, (B) Whole-body percentage fat assessed by Dual-Energy X-ray Absorptiometry, and (C, D) Soleus (SOL), tibialis anterior (TA) composition-independent IMCL concentration assessed from the ^1^H MRS of methyl protons. The relationship of soleus, tibialis anterior composition-independent IMCL concentration with VO_2max_ in a subset of participants who underwent VO_2max_ testing (controls: grey circles, n=24; athletes: white triangles, n= 14) is shown in (E, F). This was only significant when controls and athletes were combined, soleus (r = -0.52, p = 0.001) and tibialis anterior (r = -0.42, p = 0.009). * p<0.05, ** p<0.01, *** p <0.001. (A-D) tested by ANOVA and Games-Howell post hoc analysis, (E-F) by Pearson’s correlation coefficient.

**Table 1.** Characteristics of the participants, and ‘conventional’ muscle lipid estimates

|  | **Female**<br>**LD**<br>**n = 16** | **Female**<br>**controls**<br>**n = 41** | **Female**<br>**athletes**<br>**n = 14** | **P value**<br>**ANOVA** | **P value**<br>**LD-**<br>**controls** | **P value**<br>**LD-**<br>**athletes** | **P value**<br>**controls**<br>**-athletes** |
| --- | --- | --- | --- | --- | --- | --- | --- |
| Age, yr | 38.9±3.9 | 35.5±2.0 | 36.5±2.9 | 0.694 |  |  |  |
| BMI, kg.m<sup>-2</sup> | 24.2±0.7 | 24.4±0.6 | 20.3±0.6 | **<0.001** | 0.999 | **0.001** | **<0.001** |
| Mass, kg | 66.1±2.8 | 65.2±2.3 | 54.6±1.3 | **0.013** | 0.916 | **0.006** | **0.001** |
| Fat mass, kg | 10.6±1.4 | 23.1±1.6 | 11.6±1.1 | **<0.001** | **0.001** | 0.631 | **<0.001** |
| FFM, kg | 55.4±1.7 | 42.1±0.9 | 43.0±1.2 | **<0.001** | **<0.001** | **<0.001** | 0.705 |
| Triglyceride, mmol/l | 3.54±0.67 | 0.92±0.07<sup>a</sup> | 0.84±0.08 | **<0.001** | **<0.001** | **<0.001** | 0.920 |
| HDL-cholesterol, mmol/l | 0.97±0.71 | 1.64±0.06<sup>a</sup> | 2.20±0.12 | **<0.001** | **<0.001** | **<0.001** | **0.001** |
| Glucose, mmol/l | 6.00±0.58 | 4.56±0.06<sup>a</sup> | 4.46±0.13 | **<0.001** | 0.063 | **0.002** | 0.764 |
| Insulin, pmol/l | 145.6±22.0 | 38.6±4.2<sup>a</sup> | 22.6±4.7 | **<0.001** | **<0.001** | **<0.001** | **0.025** |
| HbA1c, mmol/mol | 48.3±4.2<sup>b</sup> | ND | 34.9±0.7 | ND |  | **0.008** |  |
| VO<sub>2max</sub>, ml.kg<sup>-1</sup>.min<sup>-1</sup> | ND | 36.1±1.5<sup>c</sup> | 46.9±1.4 | ND |  |  | **<0.001** |
| ***Conventional IMCL, EMCL, expressed as CH<sub>2</sub>/water-calc in units of %*** |  |  |  |  |  |  |  |
| SOL IMCL | 1.90±0.21 | 1.22±0.07 | 0.79±0.10 | **<0.001** | **0.027** | **<0.001** | **0.007** |
| TA IMCL | 0.89±0.16<sup>d</sup> | 0.61±0.04 | 0.45±0.04 | **0.035** | 0.562 | 0.115 | 0.078 |
| SOL EMCL | 2.05±0.34 | 2.22±0.19 | 1.81±0.20 | 0.493 |  |  |  |
| TA EMCL | 1.43±0.28<sup>d</sup> | 2.30±0.19 | 1.24±0.15 | **0.002** | 0.163 | 0.785 | **0.005** |

Non-normally distributed variables were log-transformed prior to performing ANOVA and Games-Howell post hoc analysis; bold P values are statistically significant. Data presented are mean ± SEM. To convert insulin pmol/l to µU/ml divide by 6.945. FFM, fat free mass; ND, not done; IMCL, intramyocellular lipid; EMCL, extramyocellular lipid; SOL, soleus; TA, tibialis anterior; CH<sub>2</sub>, methylene protons resonating at 1.3 ppm quantified as a percentage of the uncorrected calculated water resonance.
<sup>a</sup> n = 38, <sup>b</sup> n = 13, <sup>c</sup> n = 24, <sup>d</sup> n = 12.

### ^1^H MRS analysis of intramyocellular TG concentration

In the soleus muscle intramyocellular TG concentrations derived from the IMCL CH_3_ peak (0.9 ppm) (composition-independent IMCL concentrations) were not significantly increased (p = 0.477) in the LD patients compared to controls, but were higher compared to the lean athletes (p = 0.003) (Figure 3C). In the more glycolytic tibialis anterior muscle, composition-independent IMCL concentrations were similar in all three groups (Figure 3D). We also observed linear inverse correlations of VO_2max_ and IMCL concentration in the subset of controls who underwent VO_2max_ testing and athletes together (Figures 3E & F). Soleus IMCL was significantly lower in the athletes compared to the controls (p = 0.004, Figure 3C) and this remained significant (p = 0.025) compared to a subset of the controls matched for percentage body fat (body fat = 21.5±2.3 %, n = 10; vs athletes 21.1±1.8 %, n = 14; see supplementary figure (24)).

The conventional estimate of IMCL concentration using the CH_2_ resonance uncorrected for composition, CH_2_/water, showed similar trends to the composition-independent estimate using the CH_3_ peak with the exception that the LD patients’ soleus IMCL CH_2_ was significantly increased compared to controls (Table 1). This could be regarded as an artefact of the effects of compositional differences, which we consider next.

### ^1^H MRS analysis of intramyocellular lipid composition

IMCL had a significantly higher saturation index (CH_2_:CH_3_) in both muscles of the lipodystrophic patients compared to controls (soleus p = 0.008, tibialis anterior p = 0.024), but not athletes (Figures 4A & B). In the control group, smaller IMCL pools were associated with a higher saturation index, as shown by the linear regression line (dotted line in Figures 4C & D) having a gradient (ΔCH_2_/ΔCH_3_) < the mean CH_2_:CH_3_ (e.g. gradient soleus = 6.7 vs mean CH_2_:CH_3_ = 8.8; gradient tibialis anterior = 4.2 vs mean CH_2_:CH_3_ = 6.0). This phenomenon appeared to be independent of insulin sensitivity in the controls (Table 2: there was no relation of HOMA-IR with IMCL concentration). To generate a pathophysiologically meaningful measure of composition that is independent of IMCL quantity, the vertical (CH_2_) deviation from this regression line was measured and taken as a marker of the saturation of the pool that is adjusted for quantity, which we term the adjusted saturation index (CH_2_:CH_3adj_). This adjusted compositional marker was significantly higher in lipodystrophic patients compared to athletes (soleus p = 0.001, tibialis anterior p = 0.046) and also compared to controls (soleus p = 0.003) with a tendency in the tibialis anterior which just falls short of conventional statistical significance (p = 0.06) (Figures 4E & F).

**Figure 4.**
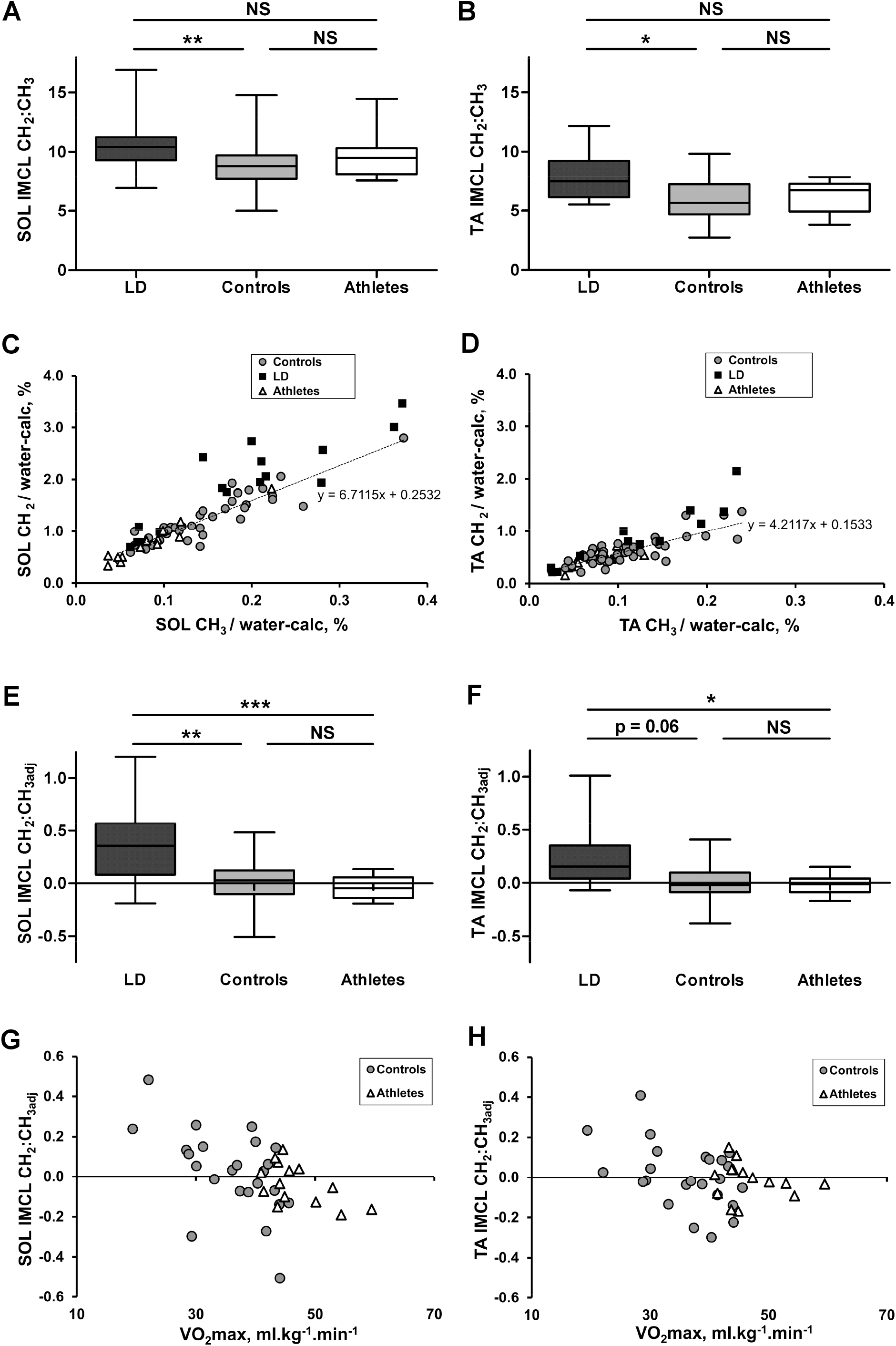
^1^H MRS measures of IMCL composition in lipodystrophic patients (black), controls (grey), and athletes (white). (A, B) Box and whisker plots of soleus (SOL) and tibialis anterior (TA) IMCL compositional saturation index (CH_2_:CH_3_ ratio). (C, D) Soleus, tibialis anterior IMCL CH_2_ and CH_3_ components. The values are expressed relative to the calculated water signal (water-calc), as described in ‘Methods’. The dotted line represents the linear regression line of the control data points. (E, F) Box and whisker plots of soleus and tibialis anterior IMCL compositional saturation index adjusted for quantity. (G, H) Relation of soleus, tibialis anterior IMCL compositional adjusted saturation index with VO_2max_ in the subset of participants who underwent VO_2max_ testing (controls: grey circles, n=24; athletes: white triangles, n= 14). Controls alone, significant correlation in the soleus (r = -0.546, p = 0.006) and tibialis anterior (r = -0.453, p = 0.026), athletes alone in the soleus (r = -0.558, p = 0.038), and with controls and athletes combined in soleus (r = -0.520, p = 0.001) and tibialis anterior (r = -0.362, p = 0.025). * p<0.05, ** p<0.01, *** p <0.001. (A-B) and (E-F) tested by ANOVA and Games-Howell post hoc analysis, (G-H) by Pearson’s correlation coefficient.

**Table 2.** Correlation coefficients of whole-body insulin resistance with IMCL

| IMCL measure | HOMA-IR |  |  |
| --- | --- | --- | --- |
|  | Controls | Controls and LD | Controls, LD and athletes |
|  | n = 38 | n = 54 <sup>a</sup> | n = 68 <sup>b</sup> |
| **Soleus** |  |  |  |
| Concentration (CH<sub>3</sub>) | -0.016 | 0.149 | 0.312* |
| Concentration and composition (CH<sub>2</sub>) | 0.153 | 0.394** | 0.477*** |
| Composition (CH<sub>2</sub>:CH<sub>3</sub>) | 0.217 | 0.439*** | 0.241* |
| Composition adjusted for quantity (CH<sub>2</sub>:CH<sub>3adj</sub>) | 0.320* | 0.583*** | 0.532*** |
| **Tibialis Anterior** |  |  |  |
| Concentration (CH<sub>3</sub>) | 0.200 | 0.205 | 0.258* |
| Concentration and composition (CH<sub>2</sub>) | 0.224 | 0.338* | 0.344** |
| Composition (CH<sub>2</sub>:CH<sub>3</sub>) | 0.100 | 0.337* | 0.202 |
| Composition adjusted for quantity (CH<sub>2</sub>:CH<sub>3adj</sub>) | 0.218 | 0.445*** | 0.364** |

\* p < 0.05, \*\* p < 0.01, \*\*\* p ≤ 0.001
Tibialis anterior has <sup>a</sup> n = 50 and <sup>b</sup> n = 64.
HOMA-IR, Homeostasis Model Assessment of Insulin Resistance; IMCL, intramyocellular lipid; CH<sub>3</sub>, methyl protons resonating at 0.9 ppm; CH<sub>2</sub>, methylene protons resonating at 1.3 ppm; CH<sub>2</sub>:CH<sub>3</sub>, compositional saturation index calculated as the ratio of CH<sub>2</sub> to CH<sub>3</sub> resonances; CH<sub>2</sub>:CH<sub>3adj</sub>, CH<sub>2</sub>:CH<sub>3</sub> saturation index adjusted for lipid quantity.

Unlike the uncorrected measure of composition, this adjusted composition also had a significant relation to VO_2max_ (Figures 4G & H) within the control subset alone, athletes alone (soleus), and control and athletes combined (statistics are given in the figure legend), such that fitter individuals had less saturated IMCL for the same absolute quantity of IMCL. VO_2max_ significantly correlated with HOMA-IR in the control subset (r = -0.59, p = 0.003, n = 23) and in controls and athletes together (r = -0.53, p = 0.001, n = 37). Table 2 shows relations of IMCL concentration and composition with insulin sensitivity.

The saturation index was higher in the soleus muscle compared with the tibialis anterior in all 3 groups. This was still the case in the two EMCL-deficient AGLD patients (P1 soleus CH_2_:CH_3_ = 9.5, TA CH_2_:CH_3_ = 6.3; P2 soleus CH_2_:CH_3_ = 11.2, TA CH_2_:CH_3_ = 7.1)

## Discussion

Using a recently validated ^1^H magnetic resonance spectroscopy method we have compared a compositional saturation index (CH_2_:CH_3_ ratio) of intramyocellular lipid in the soleus and tibialis anterior muscles of female insulin-resistant lipodystrophic patients with that of age- and gender-matched athletes and healthy controls, and shown it to be significantly higher in both muscles compared with controls, but not athletes. The finding that smaller IMCL pools in the control group had a relatively higher saturation index than larger pools irrespective of insulin sensitivity could possibly explain why the athletes studied here, who had small IMCL pools, had a statistically similar composition to patients. This observed concentration – composition relation seems physiologically plausible given that more unsaturated and shorter-chain FA are preferentially mobilized (17). We hypothesize that a person may ‘move’ along a line such as this while performing daily activities, and that a measurement of deviation from this relationship may therefore be a more sensitive and specific measure of muscle metabolic physiology or pathophysiology. To take this concentration-composition dependence into consideration we adjusted the compositional saturation index for concentration (CH_2_:CH_3adj_), and this marker was able to distinguish between athletes and lipodystrophic patients in both muscles. The strong inverse relation of CH_2_:CH_3adj_ with VO_2max_ indicates that fitness is associated with relatively lower saturation of IMCL, although overall athletes and controls were not statistically different; this is similar to results from a biopsy study (28) that demonstrated a comparable percentage saturated IMCL in controls and athletes.

By our measure of composition-independent IMCL (using the CH_3_ instead of the uncorrected CH_2_ resonance) we found that the lipodystrophic patients’ IMCL concentration was not significantly higher in either muscle compared with age-, gender- and BMI-matched controls, but was raised in the soleus relative to athletes. There are few reports of IMCL in lipodystrophic patients, but our findings are in agreement with Peterson et al. (29) who, in 3 patients with generalized lipodystrophy (2 congenital, 1 acquired), found similar soleus IMCL to 6 age-, BMI- and weight-matched controls. Calf IMCL was also lower in a case report of acquired generalized lipodystrophy (30) compared to controls, whilst a study of 4 patients with congenital generalized lipodystrophy suggested that IMCL was higher (31). Using the CH_2_ resonance, as is conventional in previous literature, we found soleus IMCL ‘content’ to be significantly higher in our LD patients compared with controls (Table 1), demonstrating, we argue, the influence of composition on measures of concentration.

### The athlete’s paradox

We found that the athletes’ IMCL was significantly lower (soleus) or similar (tibialis anterior) to controls. Although this appears to contradict well-known literature reports of an athlete’s paradox using both biopsy methods (13,28,32–34) and ^1^H MRS (35,36), this finding is in agreement with literature that reports no such paradox compared to old or young controls (37), obese individuals (38), or in certain fibre types (14). It is known that athletes can have a large depletion-repletion range of IMCL and it is possible that IMCL had not fully recovered since the last training session (24 – 48 hrs prior) as IMCL can still rise significantly after these intervals (39). In fact, our results demonstrate an inverse relation of IMCL content and VO_2max_ which is consistent with a study by Boesch et al. (40) where a combination of daily training at 60% VO_2peak_ with a low fat (10-15 % fat) diet depleted IMCL levels in both the vastus lateralis and tibialis anterior muscles to a consistent level that correlated with VO_2peak_, suggesting our female elite athletes were nearly ‘empty’ of IMCL; we did not control for diet in our study.

### Relationship of whole-body insulin resistance to IMCL

Within the controls, only the compositional saturation index adjusted for quantity (CH_2_:CH_3adj_) in soleus was significantly correlated with whole-body insulin resistance (Table 2). The inclusion of insulin-resistant patients increases the statistical significance of this relation and also yields associations with other composition-influenced markers, but not IMCL concentration in either muscle. In our study, the addition of athletes predictively reduced the associations with composition as their pools were small and therefore had a tendency for higher saturation index, but relations remain with measures that reflect large saturated pools (i.e. CH_2_:CH_3adj_ and CH_2_). These striking results suggest that the relative composition and not concentration relates to early stage insulin resistance.

### Limitations

Unlike previous ^1^H MRS studies that have conventionally reported IMCL concentrations using the predominant CH_2_ resonance whilst assuming a notional normal composition, here we have utilized the smaller CH_3_ resonance which has the advantage of composition-independence and, with comprehensive prior knowledge constraints including line width constraints relative to the CH_2_ resonance (22), fitted the IMCL CH_3_ resonance from the overlapping EMCL/IMCL CH_3_ signals. Due to fibre orientation differences between the soleus and tibialis anterior muscles the EMCL resonances are very slightly systematically shifted between muscle groups. Despite this potential for a systematic difference in the CH_2_:CH_3_ ratio between muscles, this ratio was still higher in the soleus compared to the tibialis anterior muscle in both EMCL-deficient AGLD patients, consistent with findings in our other participants in this study and a previous study (17) suggesting a compositional difference between muscles. Our patients, who mainly had partial forms of lipodystrophy, had overall similar quantities of EMCL to those of our controls and athletes which has helped in reducing potential inter-group influence of this overlapping resonance. This, together with the finding that the IMCL CH_2_:CH_3_ ratio had no relation to either EMCL CH_2_ or CH_3_ suggests a lack of EMCL influence in our datasets; however, it is possible that in other insulin-resistant cohorts large overlapping EMCL resonances may be a confounding factor.

### Summary

Use of our recently validated and potentially widely applicable ^1^H MRS approach to determine both the IMCL composition and concentration independent of composition within the soleus and tibialis anterior muscles of female individuals covering a wide range of insulin sensitivities has revealed that the IMCL composition saturation index adjusted for quantity is more strongly associated with whole-body insulin resistance than IMCL concentration alone. Differences in associations of insulin resistance with IMCL concentration when using the CH_3_ and conventional CH_2_ peaks for quantification highlights the need for awareness of the potential influence of composition on previously-reported ^1^H MRS measures of concentration.

Our finding of a strong relationship between of VO_2max_ and relatively unsaturated IMCL pools in controls and athletes points to a role of exercise in decreasing the amount of saturated fat within the IMCL store, and suggests that the composition would be able to distinguish athletes from insulin-resistant patients in cases of the athlete’s paradox. The association of insulin resistance with accumulation of saturated IMCL, even within a healthy control population, could suggest an early involvement in its pathogenesis and provide a reason why combined exercise and diet are effective therapeutic options in the early stages of insulin resistance.

## Acknowledgments

We thank all the participants, staff at both the NIHR Cambridge Clinical Research Facility and the Wolfson Brain Imaging Centre (WBIC). We acknowledge the NIHR Core Biochemistry Assay Laboratory, Cambridge Biomedical Research Centre, UK, for providing the insulin analysis. We thank Wiktor Olszowy (WBIC) for statistical advice. This research was supported by grants from the Clinical Research Infrastructure Grant, UK NIHR Cambridge Biomedical Research Centre, and the UK Medical Research Council Centre for Obesity and Related Metabolic Diseases. DBS is supported by the Wellcome Trust (107064) and AS by the NIHR via an award to the NIHR Cambridge Clinical Research Facility. The views expressed in this manuscript are those of the authors and not necessarily those of the NHS, the NIHR or the Department of Health and Social Care.

